# Predicting response to motor therapy in chronic stroke patients using Machine Learning

**DOI:** 10.1101/457416

**Authors:** Ceren Tozlu, Dylan Edwards, Aaron Boes, Douglas Labar, K. Zoe Tsagaris, Joshua Silverstein, Heather Pepper Lane, Mert R. Sabuncu, Charles Liu, Amy Kuceyeski

## Abstract

Accurate predictions of motor improvement resulting from intensive therapy in chronic stroke patients is a difficult task for clinicians, but is key in prescribing appropriate therapeutic strategies. Statistical methods, including machine learning, are a highly promising avenue with which to improve prediction accuracy in clinical practice. The first main objective of this study was to use machine learning methods to predict a chronic stroke individual’s motor function improvement after 6 weeks of intervention using pre-intervention demographic, clinical, neurophysiological and imaging data. The second main objective was to identify which data elements were most important in predicting chronic stroke patients’ impairment after 6 weeks of intervention. Data from one hundred and two patients (Female: 31%, age 61±11 years) who suffered first ischemic stroke 3-12 months prior were included in this study. After enrollment, patients underwent 6 weeks of intensive motor and transcranial magnetic stimulation therapy. Age, gender, handedness, time since stroke, pre-intervention Fugl-Meyer Assessment, stroke lateralization, the difference in motor threshold between the unaffected and affected hemispheres, absence or presence of motor evoked potential in the affected hemisphere and various imaging metrics were used as predictors of post-intervention Fugl-Meyer Assessment. Five machine learning methods, including Elastic-Net, Support Vector Machines, Artificial Neural Networks, Classification and Regression Trees, and Random Forest, were used to predict post-intervention Fugl-Meyer Assessment based on either demographic, clinical and neurophysiological data alone or in combination with the imaging metrics. Cross-validated R-squared and root of mean squared error were used to assess the prediction accuracy and compare the performance of methods. Elastic-Net performed significantly better than the other methods for the model containing pre-intervention Fugl-Meyer Assessment, demographic, clinical and neurophysiological data as predictors of post-intervention Fugl-Meyer Assessment (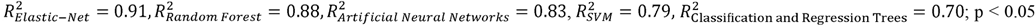). Pre-intervention Fugl-Meyer Assessment and difference in motor threshold between affected and unaffected hemispheres were commonly found as the strongest two predictors in the clinical model. The difference in motor threshold had greater importance than the absence or presence of motor evoked potential in the affected hemisphere. The various imaging metrics, including lesion overlap with the spinal cord, largely did not improve the model performance. The approach implemented here may enable clinicians to more accurately predict a chronic stroke patient’s individual response to intervention. The predictive models used in this study could assist clinicians in making treatment decisions and improve the accuracy of prognosis in chronic stroke patients.

## Introduction

Stroke is one of the most common diseases causing the functional impairment worldwide and the most common cause of the morbidity in many developed countries (Coupar *et al.*, 2012; Karahan *et al.*, 2014). Stroke survivors frequently have residual motor impairments which impact negatively their quality of life (Dobkin, 1990). One of the difficult challenges facing clinicians is to determine whether or not a patient will benefit from a certain type of treatment, particularly in the case of chronic stroke where gains may be more incremental than in the acute to sub-acute stage. Initial clinical scores, demographic information, and imaging data have been shown to be important predictors of outcome in both acute and chronic stroke patients, yet at the individual level, this remain unsatisfactory (Stinear *et al.*, 2007; Coupar *et al.*, 2012; Burke and Cramer, 2013; Asadi *et al.*, 2014; Kim and Winstein, 2017). Two of the most important predictors of upper extremity recovery are consistently found to be the initial severity of motor impairment and neurophysiological inputs, including the motor threshold (MT) and the presence of motor evoked potentials (MEP) as assessed with transcranial magnetic stimulation (TMS) (Coupar *et al.*, 2012; Jo *et al.*, 2016). In particular, one study in a small group of acute stroke subjects showed that the combined absence or presence of the MEP and the MEP threshold inputs were predictive of recovery (Manganotti *et al.*, 2015).

Magnetic resonance imaging (MRI) based measurements are widely used in stroke for diagnosis and disease monitoring of patients (Kuceyeski *et al.*, 2014, 2015a, 2016). Previous studies have shown correlation between the location of lesions with functional recovery in acute ischemic stroke (Nazzal *et al.*, 2009; Price *et al.*, 2010) and lesion size and location in motor recovery of patients with hemiplegic stroke (Chen *et al.*, 2000). In recent years, many studies have shown the importance of the connectome, i.e. the brain’s functional and structural connections as measured with MRI, in post-stroke clinical impairment and recovery (Carter *et al.*, 2010; Grefkes and Fink, 2011; Westlake and Nagarajan, 2011; Siegel *et al.*, 2016). One study in particular showed baseline metrics of the functional connectome and changes in the functional connectome from the acute to subacute stage varied between groups with different severities of impairment (Lee *et al.*, 2017). Another study showed increased baseline functional connectivity in certain regions in stroke patients who recovered better than those that did not (Puig *et al.*, 2018). Measures of structural connectome disruption have also been found to be predictive of post-stroke impairment and motor recovery. In particular, the ratio of fractional anisotropy (rFA,; a measure of white matter integrity), of the corticospinal tract between the affected and unaffected hemispheres, were found to predict motor outcome in chronic stroke patients. Lower rFA measured at day 30 was associated with a poor motor outcome at 2 years (Puig *et al.*, 2013). Here, we investigate the additive value of MRI-based measurements of structural connectome disruption that have been shown to be predictive of baseline impairment and recovery in our previous work (Kuceyeski *et al.*, 2015a, 2016). In particular, those studies used the Network Modification (NeMo) Tool (Kuceyeski *et al.*, 2013), which quantifies each gray matter region’s white matter disconnectivity to the rest of the brain as well as the amount of disconnectivity between pairs of regions. The regional measurements, called Change in Connectivity (ChaCo) scores, represent the percent of disrupted white matter fibers connecting to a given region. The pairwise disconnectivity measures represent the change in the number of fibers connecting any given pair of regions after removing those fibers that pass through the area of lesion.

Machine learning methods are highly promising for quantitative predictions of post-stoke recovery as well as assessment of demographic or imaging variables that are important in these predictions (Wang *et al.*, 2010; Cohen *et al.*, 2011). Support vector machine (SVM) and artificial neural networks (ANN) have previously shown accurate results in predicting clinical scores in chronic stroke patients (Wang *et al.*, 2014; Kim *et al.*, 2016). SVM and tree-based models (decision tree and bagging forest) have also been used to classify patients into groups that did and did not have improvement in motor function and to identify significant predictors of these classes (Rondina *et al.*, 2016; Stinear *et al.*, 2017).

The primary aim of the present study was to predict an individual’s motor function after six weeks of intervention by applying five separate machine learning methods (Elastic-net [EN], SVM, ANN, Classification and Regression Tree [CART], and Random Forest [RF]). The Fugl-Meyer Assessment (FMA), a comprehensive and consistent clinical exam, was used to assesses motor impairment both pre- and post-intervention (Duncan *et al.*, 1983; Gladstone *et al.*, 2002). Demographic, clinical, neurophysiological and imaging data were used as model input variables. Both regression (prediction of post-intervention FMA) and classification (into two groups: either clinically meaningful improvement or not) were performed with each method. The second aim was to assess the importance of variables in predicting or classifying post-intervention FMA. The third and last aim was to compare each method’s performance based on various combinations of input variables: i) demographic, clinical and neurophysiological data, ii) demographic, clinical, neurophysiological and regional disconnectivity data, and iii) demographic, clinical, neurophysiological and pair-wise disconnectivity data.

## Materials and Methods

### Subjects and intervention

The data for this paper were extracted and further analyzed from a subset of a broader multisite intervention trial with the principal results reported elsewhere (Harvey et al, 2018; NCT02089464). Institutional Review Board approval and individual consent was obtained at each site. Forty-five patients from the Burke Neurological Institute and fifty-seven patients from Rancho Los Amigos rehabilitation center were included in the present analyses. Adult (>18years) subjects with residual hemiparesis from a first-time stroke within 3-12 months prior were enrolled in the study. The intervention comprised goal-oriented intensive arm training combined with focal or diffuse non-invasive brain stimulation, three times a week for six weeks, with the primary outcome measure being the FMA. Both intervention groups were equally effective in attaining clinically meaningful improvement as per the FMA and were pooled for analysis in the present study. The following demographic, clinical and TMS-based neurophysiological information was collected for each subject: age, sex, time since stroke, handedness, left vs. right hemisphere stroke, pre-intervention FMA, difference in MT in the unaffected versus affected hemispheres and absence or presence of a MEP in the affected hemisphere. Patients were assigned a value of 100 in the affected hemisphere for the purposes of calculating the difference in MT if they did not have an MEP in the affected hemisphere, as used previously (Manganotti *et al.*, 2015).

### Image acquisition and processing

Structural MRI was acquired prior to commencement of the intervention. Lesion masks were hand-traced on patients’ native T1 scans. T1 images and the associated lesion masks were transformed to MNI space using both linear (FLIRT) and non-linear transformation techniques (FNIRT) available in FSL (http://www.fmrib.ox.ac.uk/fsl/index.html). Lesion borders were confirmed by a neurologist (A.D.B.) who was blinded to the behavioral data, both in native space and again after transformation to MNI space. Left-hemisphere lesions were flipped on the x-axis so that all lesions across both cohorts appeared in the right hemisphere. The lesion masks were then processed through the NeMo Tool software, which estimates the amount of regional disconnectivity and disconnectivity between pairs of regions based on a database of 73 healthy control brains’ white matter connectivity maps. Regional disconnectivity measurements (ChaCo scores) were quantified as the percent of white matter streamlines passing through a lesion divided by the total number connecting to that region, while pair-wise disconnectivity measurements are given as z-scores quantifying the number of white matter fibers between pairs of regions that pass through a lesion.

### Machine Learning Methods and Statistical Analysis

Five machine learning methods were used to predict post-intervention FMA and classify patients into those that improved by a clinically meaningful amount versus those that did not: 1) EN 2) ANN 3) SVM (Hsu *et al.*, 2010), 4) CART (Breiman *et al.*, 1984) and 5) RF; see Supplementary Material for details. The minimal clinically important difference in the FMA has been reported previously to be 5.5 (Page *et al.*, 2012). Thus, the patients were classified into two groups with post-intervention FMA minus pre-FMA < 5.5 or ≥ 5.5. Pre-intervention FMA, time since stroke, age, and the difference in MT were standardized to facilitate the interpretation of the intercept in the EN model. Pearson’s correlation was calculated between pairs of continuous variables in the preliminary analysis, with a significance level of 0.05 (uncorrected).

Three models for each regression/classification method were constructed for each of three sets of input variables. First, a “clinical” model was constructed that included only demographic/clinical information (age, sex, time since stroke, handedness, left vs. right hemisphere stroke, pre-intervention FMA) and TMS-based neurophysiological data (difference in MT in the unaffected versus affected hemispheres and absence or presence of a MEP in the affected hemisphere). Next, two “imaging” models were constructed that contained the variables in the clinical model plus 1) ChaCo (regional disconnectivity) scores or 2) pair-wise disconnection scores from the NeMo Tool. Model performance was assessed using the root of mean squared error (RMSE) and R-squared, defined as 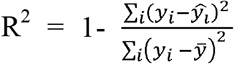 where *ŷ_i_* is the prediction of *y_i_* and *ȳ* is the mean. The performance metrics were compared across methods and input variable sets with the Kruskal-Wallis and Wilcoxon rank sum test, and considered significantly different when p < 0.05.

Each model was trained with two loops (outer and inner) of k-fold cross validation (k = 10 for both loops). The outer loop (repeated over 100 random partitions of the full dataset) provided a training set for model building and test set for model assessment. The inner loop (repeated over 10 different partitions of the training dataset only) performed grid-search to find the set of model hyperparameters that minimized the average hold-out RMSE for regression or maximized the area under Receiver Operating Characteristic curve (AUC) for classification (see the supplementary material for the intervals of the hyperparameters). A model was fitted using those optimal hyperparameters on the entire training dataset and assessed on the hold-out test set from the outer loop. At the end of the cross-validation, the mean R^2^ and RMSE in the regression analysis, and the mean AUC in the classification analysis (over each of the 100 x 10-fold test sets) were performed to assess the performance of the models. For classification, the methods provided the probability of being in the clinically meaningful recovery group (MCID≥5.5). A threshold that maximized the sensitivity of classification was used to dichotomize the results.

We considered the importance of the variables in the EN models to be the magnitude of the regression coefficient of the model averaged over all 1000 models (100 partitions of the data into 10 folds). The importance of the variables for CART models was considered to be the sum of squared error for all the splits in which a variable is used (Breiman *et al.*, 1984). In RF, each variable is randomly permuted and the difference in the new MSE and the original MSE is considered that variable’s importance (Kuhn, 2008). The importance of the variables in NNs is calculated by taking the absolute value of the input-hidden layer connection weight and dividing that by the sum of the absolute value of the input-hidden layer connection weight of all input neurons (Gevrey *et al.*, 2003). The difference in model output for the maximum and minimum values of a given variable across the subjects, while other variables are held constant at the mean across the subjects, gives The importance of the variables for SVM (Hamby, 1994).

In a post-hoc analysis, we specifically investigated the role of the neurophysiological data in improving the post-intervention FMA predictions. We compared linear models based on 1) pre-FMA only, 2) pre-FMA and difference in MT and 3) pre-FMA and the absence or presence of the MEP at the affected hemisphere. The model accuracy was compared using both the chi-squared test and weighted Akaike Information Criterion (wAIC) (Burnham and Anderson, 2004). The latter metric gives the probability that each of the possible models is the best model for the given data. The free software R (https://www.r-project.org) with version 3.4.4 was used for all statistical analyses and graphs.

### Data availability statement

The codes used to analyze the data are available upon request, but the raw data used in the analyses is proprietary and not available.

## Results

### Patient Characteristics

Table 1 shows patient demographic, clinical and neurophysiological data. Pre-intervention FMA was significantly lower than post-intervention FMA (p-value < 0.01). In an exploratory analysis for the regression models predicting post-FMA, we calculated the correlations between input variables and post-intervention FMA. The post-intervention FMA was significantly correlated with pre-intervention FMA (r= 0.954, p<0.05), time since stroke (r = −0.211, p<0.05) and the difference in MT (r = 0.626, p-value<0.05). The sign of the correlations indicates higher post-intervention FMA (better clinical score) was associated with higher pre-intervention FMA, shorter time since stroke and smaller magnitude pre-intervention inter-hemispheric difference in MT.

**Table 1.**
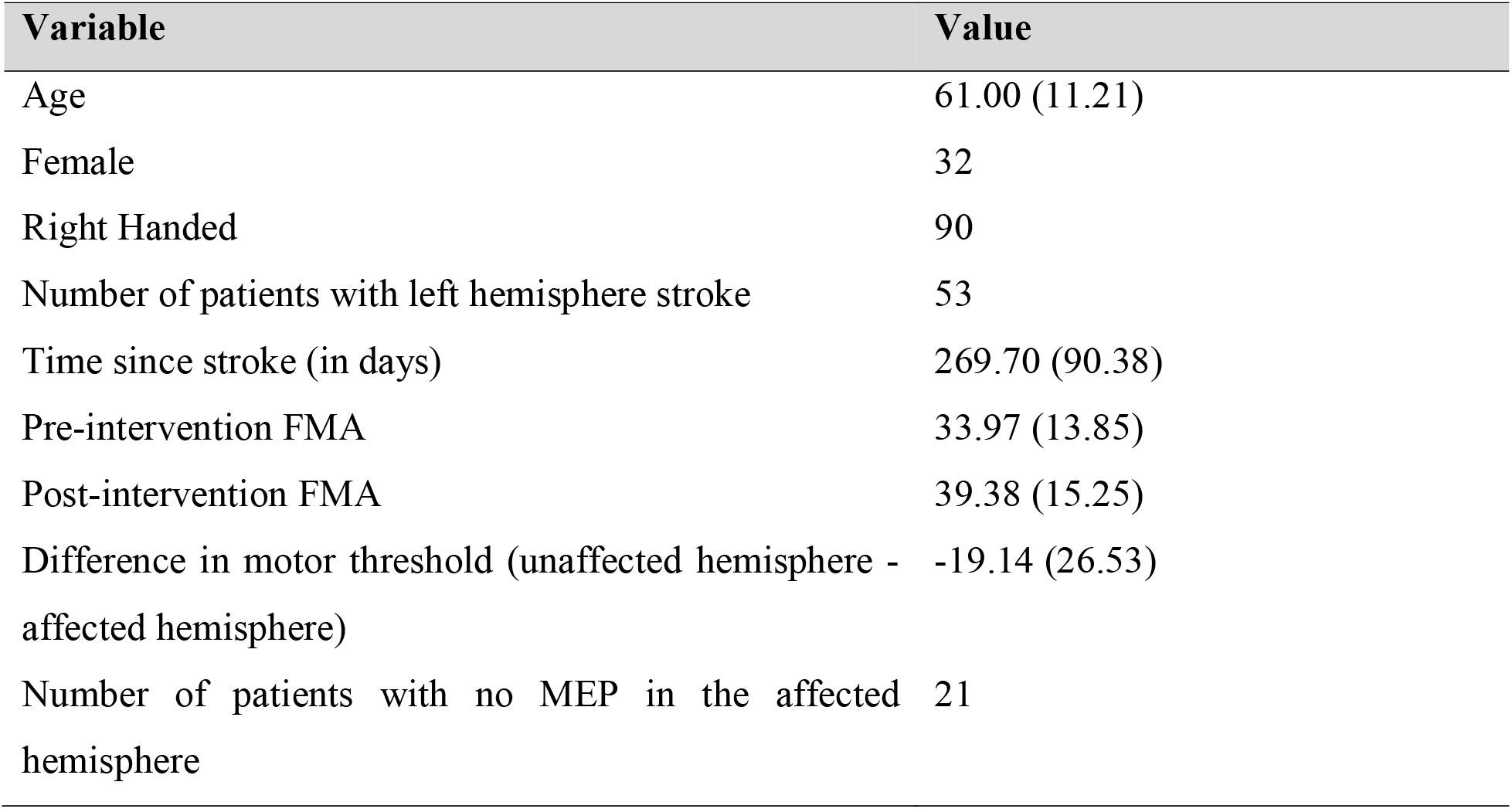
Patient demographic, clinical and neurophysiological characteristics (N = 102). Values are presented as mean (standard deviation) for the continuous variables, and as the number of patients with that characteristic for the binary variables.

### Prediction of Post-intervention FMA

Table 2 gives the R^2^ (explained variance) values for each of the five machine learning methods. EN regression showed the highest explained variance for all three sets of input variables (clinical, clinical + regional disconnectivity, and clinical + pair-wise disconnectivity). The addition of regional disconnectivity to the clinical variables resulted in less explained variance for the SVM and RF methods, and more explained variance for the ANN and CART methods. SVM and CART showed significantly higher R^2^ values with the clinical and pair-wise disconnection dataset compared to the clinical dataset only while RF and ANN had greater explained variance. The addition of both dysconnectivity measurements to the clinical model resulted in the same amount of explained variance for the EN method. Figure 1 shows violin plots of the RMSE for each of the five methods, which largely agree with the R^2^ results. Most of RMSE results were significantly different across the different sets of input variables. Only the results of SVM and CART were not significantly different for the clinical + regional disconnectivity model (p-value=0.932). The RMSE of clinical + regional disconnectivity and clinical + pair-wise disconnectivity models were significantly greater than the clinical model for all methods.

**Table 2.** The R^2^ results of the clinical, clinical + regional disconnectivity and clinical + pair-wise disconnectivity models in predicting post-intervention FMA. Values are presented as Median [1^st^ quartile, 3^rd^ quartile].

| Method | Clinical Model | Clinical + regional disconnectivity Model | Clinical + pair-wise disconnectivity Model |
| --- | --- | --- | --- |
| EN | 0.910 [0.866, 0.940] | 0.901 [0.882, 0.929] | 0.902 [0.890, 0.925] |
| RF | 0.882 [0.841, 0.917] | 0.449 [0.295, 0.510] | 0.064 [-0.148, 0.199] |
| ANN | 0.836 [0.703, 0.889] | 0.372 [-0.104, 0.650] | 0.546 [0.230,0.720] |
| SVM | 0.797 [0.729, 0.855] | 0.624 [0.448, 0.734] | 0.846 [0.744, 0.894] |
| CART | 0.703 [0.625, 0.743] | 0.790 [0.652, 0.795] | 0.823 [0.578, 0.841] |

**Figure 1.**
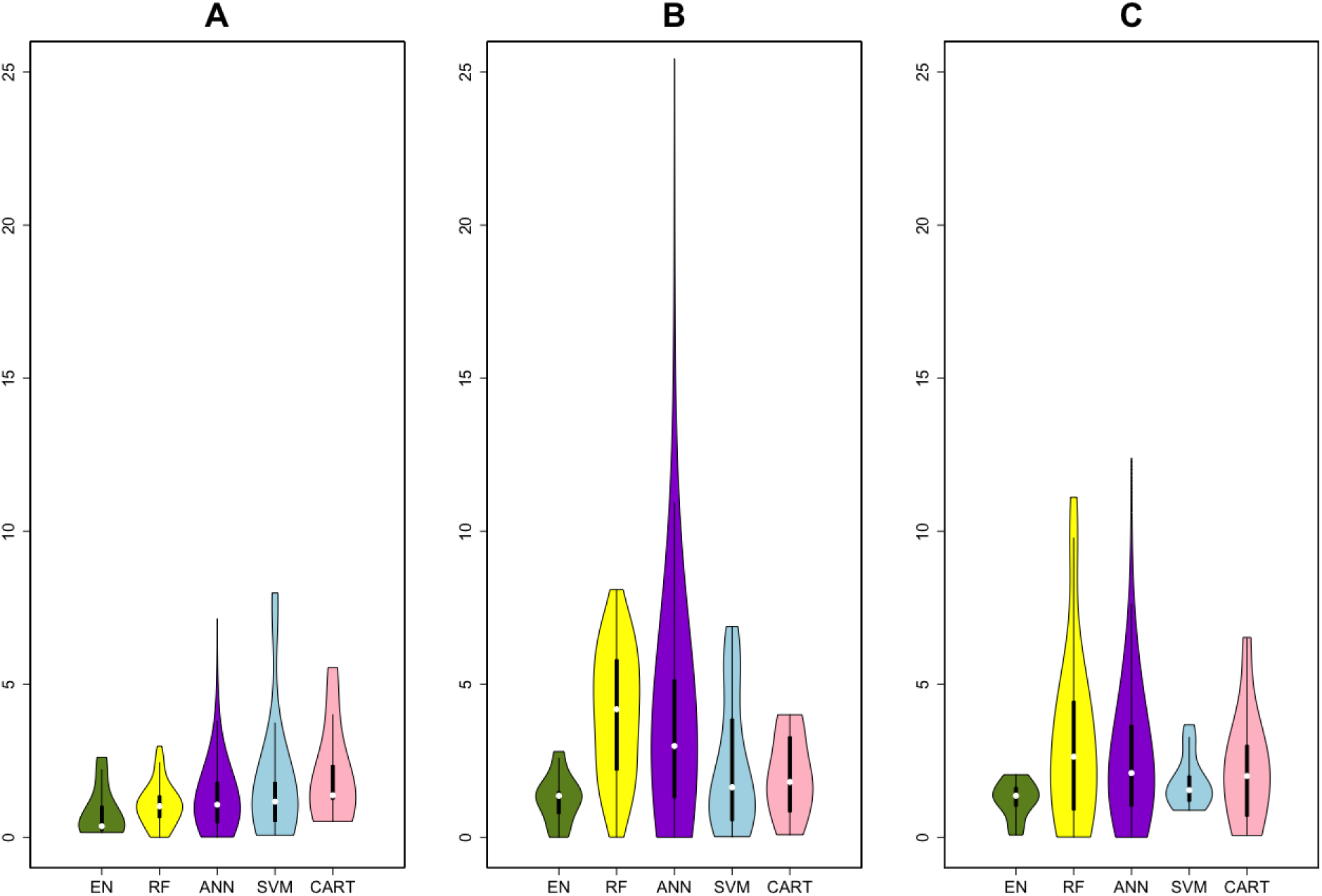
Violin plots of the RMSE for the five machine learning methods. (Elastic-Net (EN) = green, Random Forest (RF) = yellow, Artificial Neural Network (ANN) = purple, Support Vector Machines (SVM) = blue and Classification And Regression Trees (CART) = pink). The panels represent the RMSE over three sets of input variables (A) clinical, (B) clinical + regional disconnectivity and (C) clinical + pair-wise disconnectivity.

CART showed the lowest R^2^ results and EN the highest R^2^ results, with the clinical variables. Figure 2 shows the observed post-intervention FMA versus the average (+/- standard deviation) of each subject’s predictions over all 1000 models created during the cross-validation procedure. The standard deviations that were calculated by CART were larger than those that were performed by EN for most of the individuals. Also, the average predicted and observed FMA points of EN were closer to the diagonal line, and the points were linear. The predictions of CART were very dispersed for the points below the observed FMA 30.

**Figure 2.**
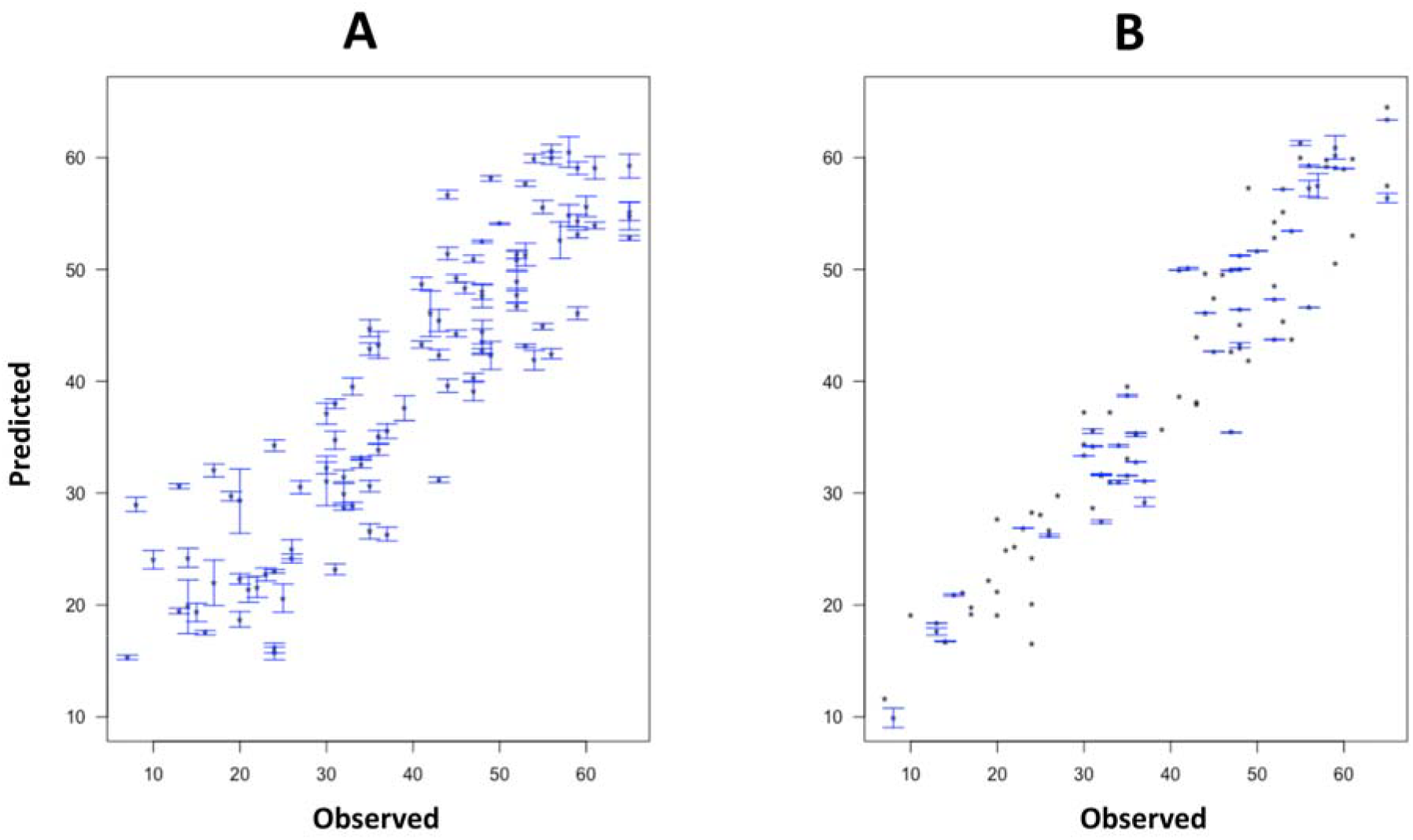
The observed versus predicted post-intervention Fugl-Meyer Assessment. Classification and Regression Tree (A), and Elastic-Net (B) models were trained on clinical input variables. Points represent the average prediction while the bars represent the standard deviation over 100 iterations in the outer loop.

Figure 3 shows bar plots of the importance of the variables for each of the five methods trained on the clinical data only. Unsurprisingly, pre-intervention FMA was the input with highest importance for all of the methods. The difference in MT between affected and unaffected hemispheres was commonly found as the next most important predictors in all the models. The absence or presence of MEP in the affected hemisphere and time since stroke were also found to be important predictors. Pre-intervention FMA and difference in MT were chosen by all 1000 models (100 iterations for 10 test data) performed with the EN. The absence or presence of MEP in the affected hemisphere, gender and right handedness were selected in more than 400 EN models (see Supplementary Table 1).

**Figure 3.**
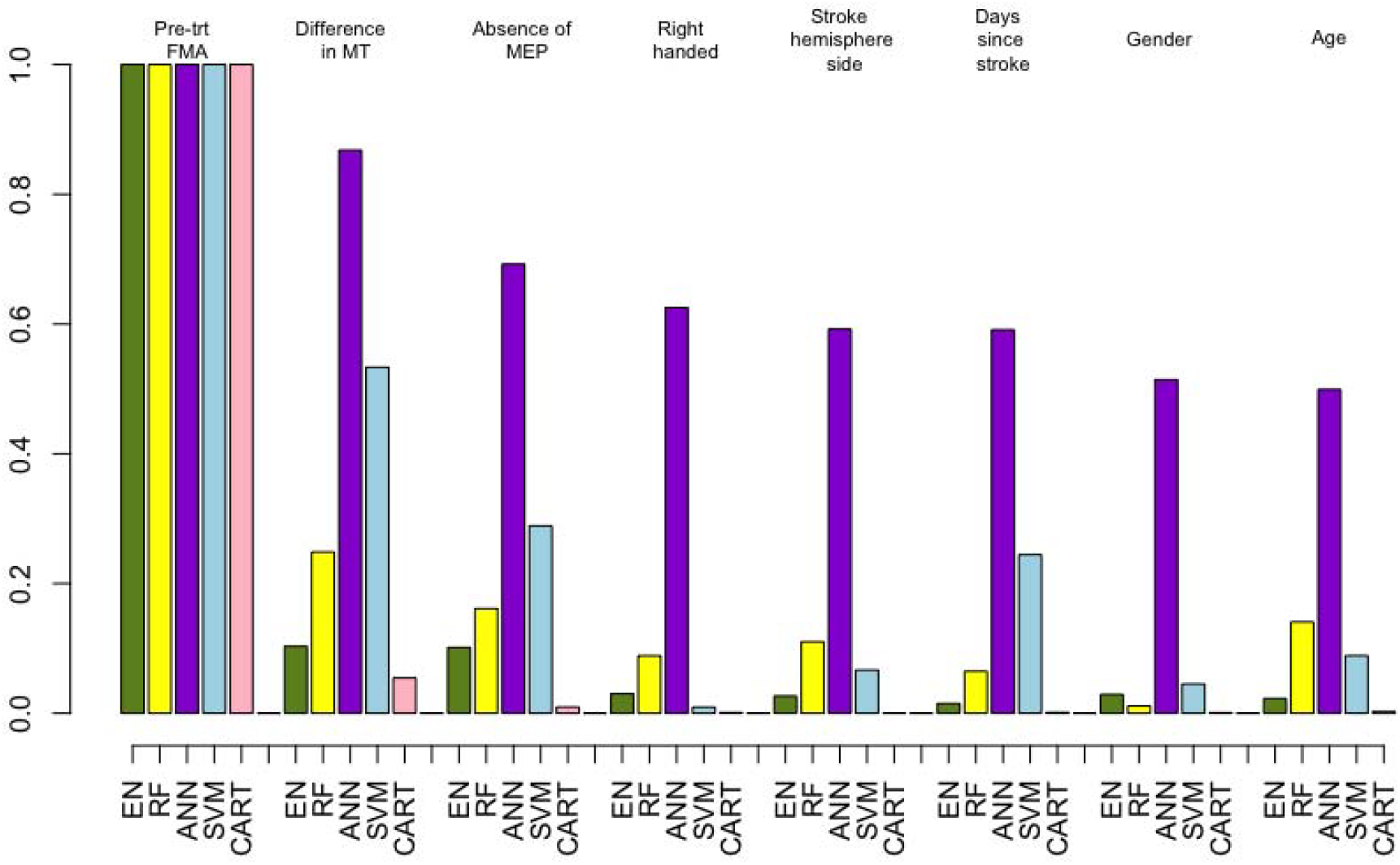
Importance of the clinical variables only (demographics, clinical and neurophysiological measures) for all five machine learning methods. (Elastic-Net (EN) = green, Random Forest (RF) = yellow, Artificial Neural Network (ANN) = purple, Support Vector Machines (SVM) = blue and Classification And Regression Trees (CART) = pink). For visualization purposes, the weights of the variables’ importance are rescaled to be relative to pre-intervention FMA.

### Classification: Good vs. Poor Recovery

Figure 4 shows violin plots of the AUC for the classification task for the 5 machine learning methods over the three sets of input variables. EN, ANN and CART gave satisfactory results, having at least one AUC greater than 0.6 over the different sets of input variables (see Supplementary Table 3). ANN and CART performed significantly better with the inclusion of the pairwise dysconnectivity data compared to the clinical model (p < 0.05). However, the performance of EN did not significantly change between the input variable sets (p-value= 0.209). The importance of the variables was also analyzed over the classification task. Only CART classification showed pre-intervention FMA as the best predictor (see Supplementary Table 2). The neurophysiological inputs were also found as important variables by some of the methods. The difference in MT was shown as the second strongest predictor by EN, RF and ANN, and the absence or presence of MEP was given as the third most important predictor by RF.

**Figure 4.**
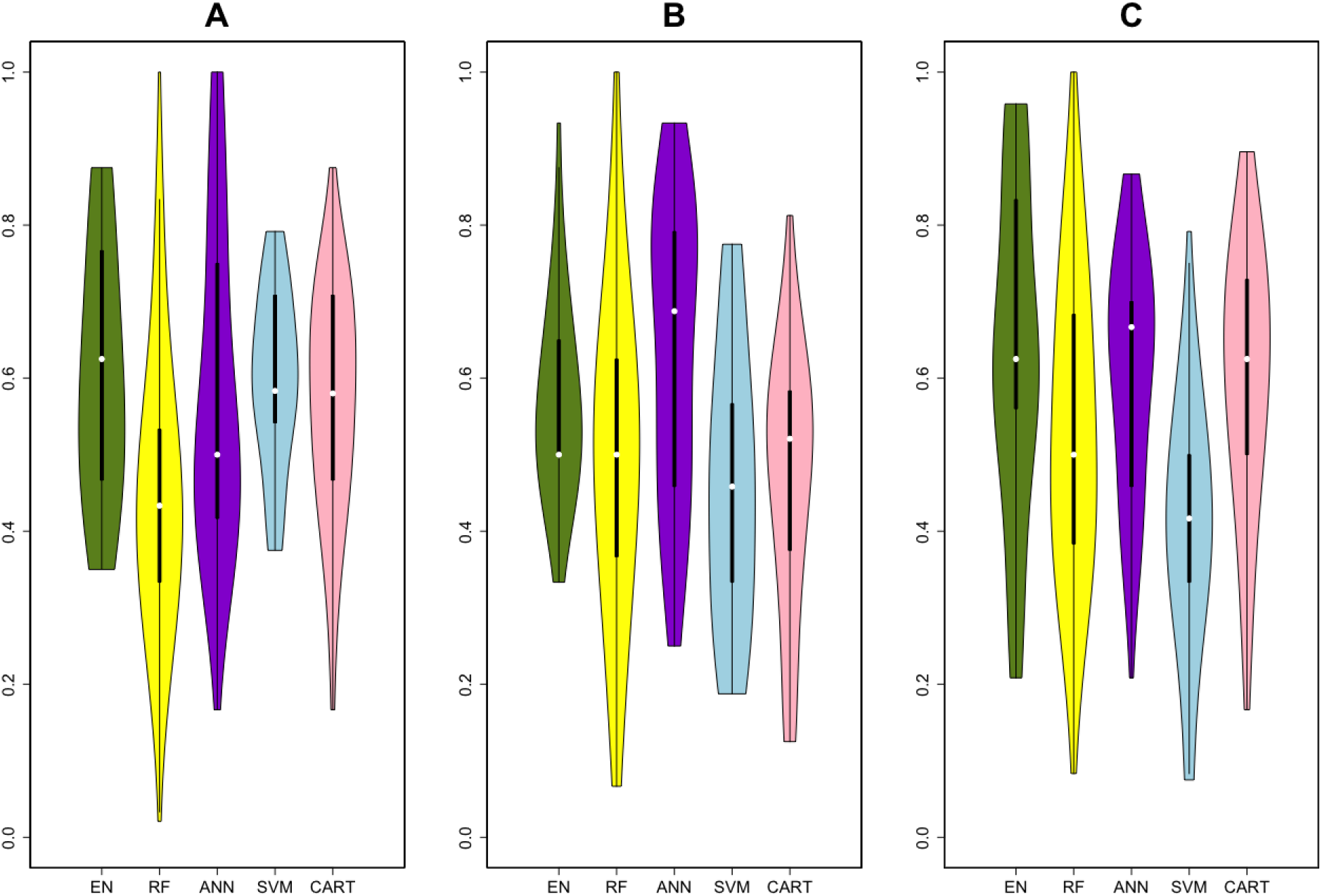
Summary of classification results (good vs. poor recovery) via violin plots of the AUC for the five machine learning methods. (Elastic-Net (EN) = green, Random Forest (RF) = yellow, Artificial Neural Network (ANN) = purple, Support Vector Machines (SVM) = blu and Classification And Regression Trees (CART) = pink) using (A) clinical, (B) clinical + regional disconnectivity and (C) clinical + pair-wise disconnectivity models.

Because of the demonstrated importance of the neurophysiological measurements in predicting post-intervention FMA, we performed the following post-hoc analysis. A linear model based on pre-intervention FMA only was compared with two linear models: one based on pre-intervention FMA and difference in MT and the other based on pre-intervention FMA and the absence or presence of the MEP at the affected hemisphere. The wAIC for the model based only on pre-intervention FMA (0.05) was much lower than for the model based on pre-intervention FMA and difference in MT (0.95). A chi-squared test showed significant improvement in the maximum likelihood when adding the difference in MT to the pre-intervention FMA only model (p-value < 0.05), while there was no change when adding the absence or presence of MEP to the pre-intervention FMA only model (p = 0.119).

## Discussion

Here we applied five machine learning methods to predict the response to intervention in a large cohort of chronic stroke patients. We predicted both a continuous measure of post-intervention motor function as well as whether or not a patient would achieve a clinically meaningful improvement in motor function. The latter is especially important if we aim to implement these types of models in a clinical setting to make decisions about patient care. This study is one of the largest to date that uses various machine learning methods on demographic, clinical, neurophysiological and imaging data to predict post-intervention motor improvements in chronic stroke. Our main findings were that EN generally out performed other methods and that neurophysiological measures derived from TMS had more influence in the prediction of recovery than other metrics, including age, time since stroke and imaging-based measures of structural disconnection. We conjecture that the EN-based models were generally better than others since it optimizes for sparsity; there were a few variables in our data that were strong predictors of recovery. Important predictors in our models, including pre-intervention motor function and TMS-derived measures, largely agree with previous findings (Hallett, 2000; Thickbroom *et al.*, 2004; Talelli *et al.*, 2006; Manganotti *et al.*, 2015; Jo *et al.*, 2016). Notably, in chronic stroke intervention outcome prediction, baseline difference in MT had greater predictive capability when added to FMA predictive models than absence or presence of MEP, and therefore should be included in prospective studies for chronic stroke outcome prediction. This may have relevance also in acute studies that typically use absence or presence of MEP in the affected hemisphere for outcome prediction, although this remains to be systemically tested in the acute recovery prediction. In fact, our post-hoc analysis showed that the difference in MT significantly improved the accuracy of prediction above and beyond a model only including pre-intervention FMA, while the absence or presence of MEP did not.

There has been increasing interest in using machine learning methods for predicting impairment and recovery stroke (Varoquaux *et al.*, 2010; Hope *et al.*, 2013; Kuceyeski *et al.*, 2015b, 2016; Stinear *et al.*, 2017). Most studies have focused on predicting recovery from the acute to sub-acute stages after stroke, when most recovery gains are made (Prabhakaran *et al.*, 2008; Stinear, 2017). One such study used ANN and SVM to make accurate predictions of acute post-stroke outcome after the intra-arterial therapy using demographic and comorbidity information (Asadi *et al.*, 2014). Rehme et al (Rehme *et al.*, 2015) used SVM method to classify patients with respect to acute post-stroke motor impairment based on the resting-state functional MRI with a classification rate of over 80%. Another study used CART analysis to classify post-stroke patients’ upper extremity motor recovery into four classes from clinical, imaging and neurophysiological data (Stinear *et al.*, 2017). They found that the absence or presence of a MEP, acute motor functionality and age were most important in predicting recovery. They largely found that imaging metrics did not significantly improve accuracy, which agrees with our current findings. However, their classification accuracy was generally around 70%, which is higher than our study’s classification results. This could be due to many differences between our studies, the most likely of which is that our predictions were for changes in chronic stroke due to intervention and not in predicting acute recovery. The former is a much more difficult task, as the changes in the chronic stage of stroke are most often not as large and thus have a lower signal-to-noise ratio. However, attempts have been made to predict response of chronic stroke subjects to treatment or interventions, although most are in moderate sample sizes or are correlation based and do not utilize machine learning techniques (Gauthier *et al.*, 2012; Ramos-Murguialday *et al.*, 2013). One study in particular found that measures of functional and structural connectivity were important in predicting motor gains from therapy in chronic stroke; however their overall predictive accuracy was moderate (R^2^ = 0.44) and their sample size relatively small (N = 29)(Abdelnour, Mueller, and Raj 2014).

Our imaging measures were based on the NeMo Tool’s estimates of structural disconnection that resulted from the lesion mask, not the lesion masks themselves. One study in 30 chronic stroke patients used SVM applied to voxel-wise probability lesion masks to classify patients into those with the most recovery versus the least recovery, and found an AUC of around 0.73 (Rondina *et al.*, 2017). We attempted to use our binary lesion masks in a similar fashion but only had an AUC of around 0.5 (results not shown). However, the group definitions in (Rondina *et al.*, 2017) were somewhat arbitrary (40% best recovered versus 40% worst recovered), they used probabilistic lesion masks, their sample size about a third of the size of this study and they focused on predicting recovery from the acute to chronic stage.

One limitation of this study is that we had only the patient’s structural T1 scans. It may be more informative to have access to stroke individual’s diffusion or functional MRI, which have been shown to be important for extracting biomarkers that can predict impairment and recovery after stroke (Stinear *et al.*, 2007; Siegel *et al.*, 2016; Puig *et al.*, 2018). One recent study showed increased baseline functional connectivity in certain regions in stroke patients who recovered better than those that did not (Puig *et al.*, 2018). Other studies have shown biomarkers of structural white matter integrity, particularly in the motor tracts, are predictive of recovery (Stinear *et al.*, 2007; Lindenberg *et al.*, 2012; Rüber *et al.*, 2012). Future studies in this cohort will focus on collecting multi-modal imaging data at baseline and post-intervention to make better predictions of response to treatment as well as detect recovery-relevant changes that may shed light on neurological mechanisms of motor improvements. Another limitation of this study was the sample size; while relatively large for studies of this nature, there are machine learning techniques (such as deep learning) that may improve accuracy of predictions but may only be implemented on much larger sets of data.

## Conclusions

In summary, this study provided a thorough comparison of various machine learning methods in predicting the response of chronic stroke patients to motor therapy. We compared models trained on various sets of input variables, including demographic, clinical, neurophysiological and imaging-based (disconnectivity) data. All machine learning methods gave highly promising results in predicting post-intervention FMA using demographic, clinical and neurophysiological data, in particular pre-intervention FMA and difference in MT in the affected versus unaffected hemisphere. Thorough validation of these types of machine learning methods using larger, multi-modal sets of data is needed. If successful, these models will provide for clinicians a valuable tool that can improve the accuracy of prognoses and response to treatment, which in turn can assist in developing personalized therapeutic plans.

## Acknowledgements

C.T. carried out the statistical analyses, drafted and wrote the article.

D.E. performed study oversight, data interpretation, manuscript development

A.B. checked the data, commented and reviewed the article.

K. Z. T., CL and H. P. L. organized, checked, and managed the data collection, commented and reviewed the article.

J.S. performed neurophysiology data acquisition, processing and interpretation

M.S. supervised the analysis and reviewed the article.

A.K. organised the study, commented and reviewed the article.

## Abbreviations

ANN: Artificial Neural Networks
AUC: Area Under Receiver Operating Characteristic Curve
CART: Classification and Regression Trees
ChaCo: Change in Connectivity
EN: Elastic-Net
FMA: Fugl-Meyer Assessment
GI: Gini Index
MCID: Minimal Clinically Important Difference
MEP: Motor Evoked Potential
MSE: Mean Squared Error
MT: Motor Threshold
NeMo Tool: Network Modification Tool
RMSE: Root of Mean Squared Error
RF: Random Forest
SVM: Support Vector Machine
TEP: Transcranial magnetic stimulation evoked potential
TMS: Transcranial magnetic stimulation
wAIC: Weighted Akaike Information Criterion

## References

Abdelnour F, Mueller S, Raj A. Relating cortical atrophy in temporal lobe epilepsy with graph diffusion-based network models. PLoS Comput. Biol 2014; 90: 335–347.

Asadi H, Dowling R, Yan B, Mitchell P. Machine learning for outcome prediction of acute ischemic stroke post intra-arterial therapy. PLoS One 2014; 9: 14–19.

Breiman L, Friedman J, J. Stone C, Olshen RA. Classification Algorithms and Regression Trees [Internet]. 1984. Available from: https://rafalab.github.io/pages/649/section-11.pdf

Burke E, Cramer SC. Biomarkers and predictors of restorative therapy effects after stroke. Curr Neurol Neurosci Rep. 2013; 13: 1–14.

Burnham KP, Anderson DR. Multimodel inference: Understanding AIC and BIC in model selection. Sociol. Methods Res. 2004; 33: 261–304.

Carter AR, Astafiev S V, Lang CE, Connor LT, Strube MJ, Pope DLW, et al. Resting Inter-hemispheric fMRI Connectiviyt Predicts Performance after Stroke. Ann. Neurol. 2010; 67: 365–375.

Chen CL, Tang FT, Chen HC, Chung CY, Wong MK. Brain lesion size and location: effects on motor recovery and functional outcome in stroke patients. Arch. Phys. Med. Rehabil. 2000; 81: 447–52.

Cohen JR, Asarnow RF, Sabb FW, Bilder RM, Bookheimer SY, Knowlton BJ, et al. Decoding continuous variables from neuroimaging data: Basic and clinical applications. Front. Neurosci. 2011; 5: 1–12.

Coupar F, Pollock A, Rowe P, Weir C, Langhorne P. Predictors of upper limb recovery after stroke: A systematic review and meta-analysis. Clin. Rehabil. 2012; 26: 291–313.

Dobkin BH. Rehabilitation after stroke. N. Engl. J. Med. 1990; 352: 1677–84.

Duncan PW, Propst M, Nelson SG. Reliability of the Fugl-Meyer assessment of sensorimotor recovery following cerebrovascular accident. Phys. Ther. 1983

Gauthier L V., Taub E, Mark VW, Barghi A, Uswatte G. Atrophy of Spared Gray Matter Tissue Predicts Poorer Motor Recovery and Rehabilitation Response in Chronic Stroke. Stroke 2012; 43: 453–457.

Gevrey M, Dimopoulos I, Lek S. Review and comparison of methods to study the contribution of variables in artificial neural network models. Ecol. Model. 160 2003; 160: 249–264.

Gladstone DJ, Danells CJ, Black SE. The Fugl-Meyer Assessment of Motor Recovery after Stroke: A Critical Review of Its Measurement Properties. Am. Soc. Neurorehabilitation 2002; 16: 232–240.

Grefkes C, Fink GR. Reorganization of cerebral networks after stroke: New insights from neuroimaging with connectivity approaches. Brain 2011; 134: 1264–1276.

Hallett M. Transcranial magnetic stimulation and the human brain. Nature 2000; 406: 147–150.

Hamby DM. A review of techniques for parameter sensitivity analysis of environmental models. Environ. Monit. Assess. 1994; 32: 135–154.

Hope TMH, Seghier ML, Leff AP, Price CJ. Predicting outcome and recovery after stroke with lesions extracted from MRI images. NeuroImage Clin. 2013; 2: 424–433.

Hsu C, Chang C, Lin C. A Practical Guide to Support Vector Classification. 2010; 1: 1–16.

Jo JY, Lee A, Kim MS, Park E, Chang WH, Shin Y-I, et al. Prediction of Motor Recovery Using Quantitative Parameters of Motor Evoked Potential in Patients With Stroke. Ann. Rehabil. Med. 2016; 40: 806–815.

Karahan AY, Kucuksen S, Yilmaz H, Salli A, Gungor T, Sahin M. Effects of Rehabilitation Services on Anxiety, Depression, Care-Giving Burden and Perceived Social Support of Stroke Caregivers. Acta Medica (Hradec Kral. Czech Republic) 2014; 57: 68–72.

Kim B, Winstein C. Can Neurological Biomarkers of Brain Impairment Be Used to Predict Poststroke Motor Recovery? A Systematic Review. Neurorehabil. Neural Repair 2017; 31: 3–24.

Kim WS, Cho S, Baek D, Bang H, Paik NJ. Upper extremity functional evaluation by Fugl-Meyer assessment scoring using depth-sensing camera in hemiplegic stroke patients. PLoS One 2016; 11: 1–13.

Kuceyeski A, Kamel H, Navi BB, Raj A, Iadecola C. Predicting future brain tissue loss from white matter connectivity disruption in ischemic stroke. Stroke 2014; 45: 717–722.

Kuceyeski A, Maruta J, Relkin N, Raj A. The Network Modification (NeMo) Tool: Elucidating the Effect of White Matter Integrity Changes on Cortical and Subcortical Structural Connectivity. Brain Connect. 2013; 3: 451–463.

Kuceyeski A, Navi BB, Kamel H, Raj A, Relkin N, Toglia J, et al. Structural connectome disruption at baseline predicts 6-months post-stroke outcome. Hum. Brain Mapp. 2016; 37: 2587–2601.

Kuceyeski A, Navi BB, Kamel H, Relkin N, Villanueva M, Raj A, et al. Exploring the brain’s structural connectome: A quantitative stroke lesion-dysfunction mapping study. Hum. Brain Mapp. 2015a; 36: 2147–2160.

Kuceyeski A, Navi BB, Kamel H, Relkin N, Villanueva M, Raj A, et al. Exploring the brain’s structural connectome: a quantitative stroke lesion-dysfunction mapping study. Hum. Brain Mapp. 2015b; 36: 2147–2160.

Kuhn M. Building Predictive Models in R Using the caret Package. J. Stat. Softw. 2008; 28: 1–26.

Lee J, Park E, Lee A, Chang WH, Kim DS, Kim YH. Recovery-related indicators of motor network plasticity according to impairment severity after stroke. Eur. J. Neurol. 2017; 24: 1290–1299.

Lindenberg R, Zhu LL, Rüber T, Schlaug G. Predicting functional motor potential in chronic stroke patients using diffusion tensor imaging. Hum. Brain Mapp. 2012; 33: 1040–51.

Manganotti P, Acler M, Masiero S, Del Felice A. TMS-evoked N100 responses as a prognostic factor in acute stroke. Funct. Neurol. 2015; 30: 125–130.

Nazzal ME, Saadah MA, Saadah LM, Trebinjac SM. Acute ischemic stroke: relationship of brain lesion location & functional outcome. Disabil. Rehabil. 2009; 31: 1501–6.

Page SJ, Fulk GD, Boyne P. Clinically Important Differences for the Upper-Extremity Fugl-Meyer Scale in People With Minimal to Moderate Impairment Due to Chronic Stroke. Am. Phys. Ther. Assoc. 2012; 92: 791–798.

Prabhakaran S, Zarahn E, Riley C, Speizer A, Chong JY, Lazar RM, et al. Inter-individual Variability in the Capacity for Motor Recovery After Ischemic Stroke. Neurorehabil. Neural Repair 2008; 22: 64–71.

Price CJ, Seghier ML, Leff AP. Predicting language outcome and recovery after stroke: the PLORAS system. Nat. Rev. Neurol. 2010; 6: 202–10.

Puig J, Blasco G, Alberich-Bayarri A, Schlaug G, Deco G, Biarnes C, et al. Resting-State Functional Connectivity Magnetic Resonance Imaging and Outcome After Acute Stroke. Stroke 2018; 49: 2353–2360.

Puig J, Blasco G, Daunis-I-Estadella J, Thomalla G, Castellanos M, Figueras J, et al. Decreased corticospinal tract fractional anisotropy predicts long-term motor outcome after stroke. Stroke 2013; 44: 2016–2018.

Ramos-Murguialday A, Broetz D, Rea M, LäerL, Yilmaz Ö, Brasil FL, et al. Brain-machine interface in chronic stroke rehabilitation: A controlled study. Ann. Neurol. 2013; 74: 100–108.

Rehme AK, Volz LJ, Feis DL, Bomilcar-Focke I, Liebig T, Eickhoff SB, et al. Identifying neuroimaging markers of motor disability in acute stroke by machine learning techniques. Cereb. Cortex 2015; 25: 3046–3056.

Rondina JM, Filippone M, Girolami M, Ward NS. Decoding post-stroke motor function from structural brain imaging. NeuroImage Clin. 2016; 12: 372–380.

Rondina JM, Park CH, Ward NS. Brain regions important for recovery after severe post-stroke upper limb paresis. J. Neurol. Neurosurg. Psychiatry 2017; 88: 737–743.

Rüber T, Schlaug G, Lindenberg R. Compensatory role of the cortico-rubro-spinal tract in motor recovery after stroke. Neurology 2012; 79: 515–22.

Siegel JS, Ramsey LE, Snyder AZ, Metcalf N V., Chacko R V., Weinberger K, et al. Disruptions of network connectivity predict impairment in multiple behavioral domains after stroke. Proc. Natl. Acad. Sci. 2016; 113: 4367–4376.

Stinear CM. Prediction of motor recovery after stroke: advances in biomarkers. Lancet Neurol. 2017; 16: 826–836.

Stinear CM, Barber PA, Smale PR, Coxon JP, Fleming MK, Byblow WD. Functional potential in chronic stroke patients depends on corticospinal tract integrity. Brain 2007; 130: 170–180.

Stinear CM, Byblow WD, Ackerley SJ, Smith M-C, Borges VM, Barber PA. PREP2: A biomarker-based algorithm for predicting upper limb function after stroke [Internet]. Ann. Clin. Transl. Neurol. 2017; 4(11): 811–820.

Talelli P, Greenwood RJ, Rothwell JC. Arm function after stroke: Neurophysiological correlates and recovery mechanisms assessed by transcranial magnetic stimulation. Clin. Neurophysiol. 2006; 117: 1641–1659.

Thickbroom GW, Byrnes ML, Archer SA, Mastaglia FL. Motor outcome after subcortical stroke correlates with the degree of cortical reorganization. Clin. Neurophysiol. 2004; 115: 2144–2150.

Varoquaux G, Baronnet F, Kleinschmidt A, Fillard P, Thirion B. Detection of Brain Functional-Connectivity Difference in Post-stroke Patients Using Group-Level Covariance Modeling. Springer, Berlin, Heidelberg; 2010. p. 200–208.

Wang J, Yu L, Wang J, Guo L, Gu X, Fang Q. Automated Fugl-Meyer Assessment using SVR model. 2014 IEEE Int. Symp. Bioelectron. Bioinformatics, IEEE ISBB 2014 2014: 0–3.

Wang Y, Fan Y, Bhatt P, Davatzikos C. High-dimensional pattern regression using machine learning: From medical images to continuous clinical variables. Neuroimage 2010; 50: 1519–1535.

Westlake KP, Nagarajan SS. Functional Connectivity in Relation to Motor Performance and Recovery After Stroke. Front. Syst. Neurosci. 2011; 5: 1–12.

